# Drought and flooding resistances of maize germplasm are compared and evaluated using a multivariate analysis method

**DOI:** 10.1101/2022.08.11.503629

**Authors:** Guo Yun Wang, Shakeel Ahmad, Yong Wang, Bing Wei Wang, Jing Hua Huang, Mohammad Shah Jahan, Xun Bo Zhou, Cheng Qiao Shi

**Affiliations:** Guangxi Key Laboratory of Agro-environment and Agro-products Safety, Guangxi Colleges and Universities Key Laboratory of Crop Cultivation and Tillage, National Demonstration Center for Experimental Plant Science Education, Agricultural College of Guangxi University, Guangxi, Nanning 530004, China; Maize Research Institute, Guangxi Academy of Agricultural Sciences, Guangxi, Nanning 530007, China

**Keywords:** Maize, Drought and Flooding Stresses, Mechanism, Cultivar Verification

## Abstract

Drought and flooding stress alternately and frequently occur in Guangxi, China, and the whole world, which seriously limit maize production. Few studies focus on the different responses and evaluations of maize to drought and flooding stresses. A pot experiment with 40 varieties was conducted under well water, drought and flooding stresses. A multivariate analysis method of principal component analysis, comprehensive evaluation value, correlation analysis, stepwise regression analysis, and cluster analysis was used to evaluate the resistance of maize. Most varieties had stronger drought resistance rather than flooding resistance because of the higher antioxidant enzyme activities, osmotic adjustment substances, less reactive oxygen species, and a greater than 1.0 drought-resistance coefficient. However, there was an increment of reactive oxygen species (especially O_2_^−^), ascorbate peroxidase, peroxidase, soluble sugar, and the decrement of superoxide dismutase, catalase, soluble protein, and a lower than 1.0 of flooding-resistance coefficient of most maize varieties in flooding stress compared with well water. The superoxide dismutase, peroxidase, catalase, ascorbate peroxidase, proline, soluble sugar and protein, plant height, leaf area/plant, and stem diameter were screened out to be accurate and representative indicators to evaluate the drought and flooding resistance of maize. The study provides an insight to comprehend the different mechanisms of maize in response to drought and flooding stresses and provides a multivariate analysis method for screening the resistance of maize germplasm which could be valuable for further research and breeding of drought and flooding resistances of maize.

**One-sentence summary:** A multivariate analysis method for the screening the resistance of maize germplasm and the different physiological mechanisms of drought and flooding stresses were revealed.

## Introduction

China was the 1^st^ most affected country by flooding in the world which experienced an average of 20 flooding occurrences per year during 2000-2019 (EM-DAT2020). Meanwhile, drought marked the 2^nd^ most impactful disaster, which affected 1.4 billion people (EM-DAT2020). The current trajectory for frequency and intensity of drought and flooding are becoming more and more serious changed with the global climate (Forootan et al., 2019; He et al., 2020), which become the major abiotic stresses that limit crop growth and production (GRFC2020; Ahmad et al., 2022). Guangxi, located in the south subtropical region of China, is the main planting area for the maize double cropping system. Maize (*Zea mays* L.) in the area is largely rainfed, nevertheless, the temporal and spatial distribution of rainfall is extremely uneven as the extreme climates of flooding and drought often alternately occur especially in the seedling stage of maize in Guangxi. According to the National Bureau of Statistics of China, an annual average of 203.6 thousand ha and 453.5 thousand ha of crops area suffered from drought and flooding damages across 2010-2018 respectively. Abrupt drought-flooding alternation results in a series of widespread agriculture, forestry, livestock, etc. economic loss, outbreaks of epidemic diseases, and people death (He et al., 2020; Ahmad et al., 2021). To alleviate the damage of drought and flooding stress (called water stress) to maize, so far, there is little research basis on the drought and flooding resistant characteristics of maize varieties in Guangxi, so evaluating and screening maize for drought and flooding resistance identification indices will be great significance to resistance study and breeding of maize.

Maize is more susceptible to water stress in its vegetative stage (Huang et al., 2019; Goyal et al., 2020), particularly in the third-leaf stage (Chen et al., 2016; Shin et al., 2016; Ren et al., 2018). The chlorophyll, photosynthetic capacity, dry matter, and yield in the third-leaf stage were significantly lower than those in the jointing and tasseling stages (Tian et al., 2019). Maize at a third-leaf stage in autumn in Guangxi often suffered from the damage of abrupt drought-flooding alternation which resulted in massive production reduction. To adverse the effects of water stress, crops exhibit specific physiological changes through self-regulation in resilience to water stress. Osmotic adjustment substances including soluble sugar, protein and proline, and antioxidant enzymes comprised of superoxide dismutase (SOD), peroxidase (POD), catalase (CAT), ascorbate peroxidase (APX), as two of the most important physiological changes are increased under water stress (Nikolaeva et al., 2015; Anjum et al., 2016; Lukić et al., 2021). They all play an essential role in eliminating the damaging effects of reactive oxygen species (ROS) in plant cells in response to water stress (Huang et al., 2019; Ahmad et al., 2022). Of course, those are only two of the complicated resistant mechanism of crops in answer to water stress. However, they combined with morphological indicators can be quoted as more representative indices for the identification of drought resistance (Zou et al., 2020; Sun et al., 2021).

The drought or flooding resistant mechanism of crops is fairly complicated, and cannot be fully and accurately evaluated by a single index alone. At present, many indicators have been widely used in analyzing drought resistance (Dossa et al., 2017; Aghaie et al., 2018), but the number and efficiency of measurement indicators are relatively more and lower respectively. Meanwhile, a particular correlation is detected among many evaluation indicators like a linear correlation between yield with membership function value (Sun et al., 2021), leading to the overlapping of the information when indexes are used to evaluate the resistance of crops (Zou et al. 2020). On account of that, a multivariate analysis method is necessary to be applied to evaluate the resistance of crops’ comprehensive character indicators. Scientists have jointed the application of two or more multivariate analysis methods such as comprehensive drought-resistance coefficient, correlation, multiple linear regression analysis (Quevedo et al., 2022), principal component analysis (PCA, Wijewardana et al., 2016; Füzy et al., 2019), membership function value, comprehensive evaluation (D) value and cluster analysis to evaluate the resistance of crops (Shahrokhi et al., 2020; Yu et al., 2021). Zou et al. (2020) combined PCA, stepwise regression, and D value to screen out proline, SOD, CAT, etc, nine typical indices associated with drought resistance avoiding the one-sidedness of a single index and overlapping of the information. The relationship between crop drought tolerance and tolerance traits is also revealed through multi-methods comprehensive analysis (Füzy et al., 2019; Sun et al., 2021). A more reliable and practical multivariate analysis method will play an important role in and be able to be widely applied to evaluate and screen cultivars of crops.

At present, limited studies have been carried out to screen either the flooding tolerance or the tolerance to both stresses of drought and flooding of maize cultivars, especially in the flood-prone area. Therefore, the objectives of our study were to select drought or flooding-resistance indicators and to identify drought or flooding resistance ability as well as the differences lied in different maize varieties through a multivariate statistical analysis.

## Materials and Methods

### Plant material and water treatment

The experiment was conducted in the net room of Guangxi University, Nanning, Guangxi (latitude 22°50’N, longitude 108°17’E), China. Humid sub-tropical monsoon climate is the prevailing climate with an annual average air temperature of 21.8 °C, precipitation of 1309.7 mm and relative humidity of 79.1% in 30 years. The temperature of 33.0°C and relative humidity of 65.4% was detected during the experiment in July, 2021 (Figure 1). The soil of sandy clay loam (World Reference Base) from the field of 0–20 cm soil layer was characterized by a pH 6.8, field capacity of 30.3% (g/g, %), soil bulk density of 1.2 g/cm^3^, soil organic matter 10.4 g/kg, and available nitrogen, phosphorus and potassium 90.7 mg/kg, 83.7 mg/kg and 293.0 mg/kg, respectively.

**Figure 1.**
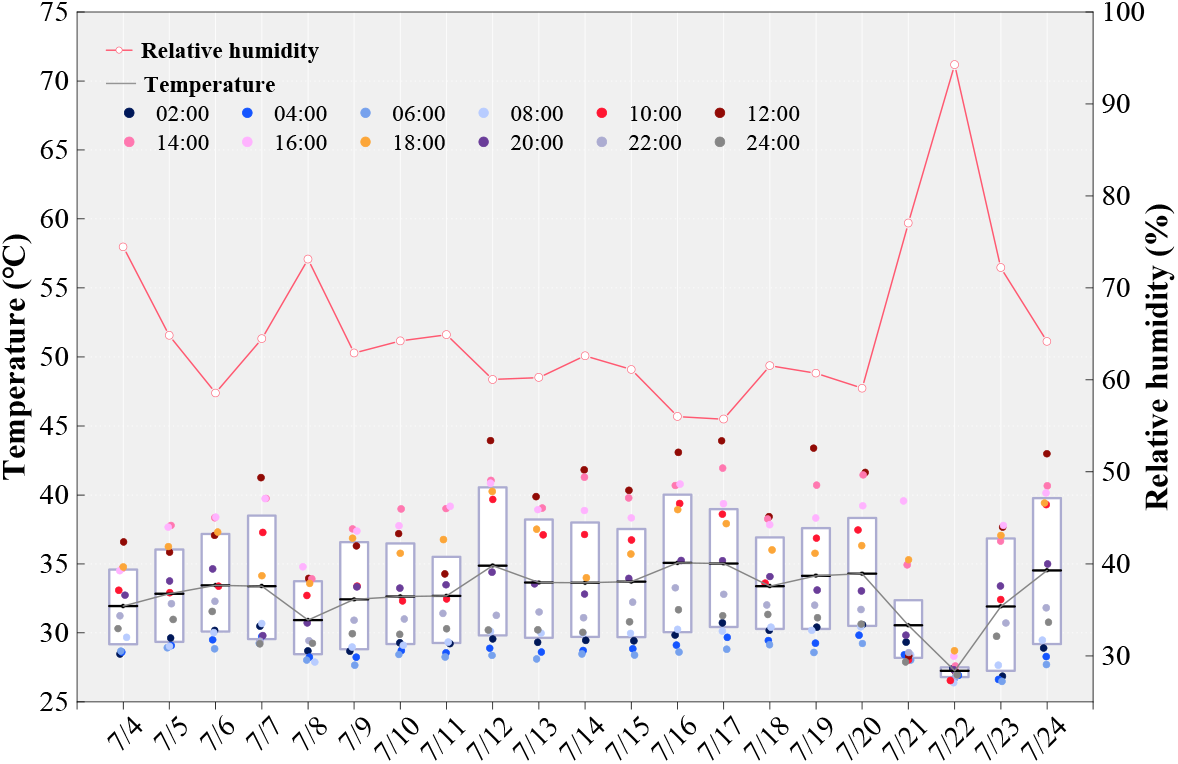
Cumulative daily air temperature and relative humidity during the experiment in July, 2021.

Split-plot experimental design with three replications was adopted in the experiment. Three water treatments including well water (CK, 70.0–75.0% field capacity), drought stress (40.0–45.0% field capacity) and flooding with a 2.0-3.0 water layer were designed as the main plot. Forty maize varieties provided by Maize Research Institute, Academy of Agricultural Sciences, Guangxi, China, was designed as split plot (Table 1). Each pot about 23.6 cm in diameter and 24.5 cm high was filled with 8.0 kg field soil of 0-20 cm that contained 3.8% soil water content and 2.3 g urea (46.2% N), 2.3 g calcium magnesium phosphate (18.0% P_2_O_5_), and 0.7 g muriate of potash (60.0% K_2_O). Ten seeds were planted in each pot on 4th July, 2021, and the seedlings were thinned to five per pot at the two-leaf stage. Water stress was treated at the three-leaf stage and then lasted for eight days, but no irrigation in drought stress was given two days before the three-leaf stage controlling the soil water content lower than 60.0% field capacity. Weighing pots every two days to control soil water content. Another big pot (25.0 cm × 29.5 cm) was set together with each pot in flooding to maintain the water layer. After eight days treated, all fully developed leaves per pot were sampled and pooled together to store at −80 °C until further analysis.

**Table 1.** The maize varieties were applied in the experiment.

| Code | Varieties | Code | Varieties | Code | Varieties | Code | Varieties |
| --- | --- | --- | --- | --- | --- | --- | --- |
| 1 | Guidan 162 | 11 | Guidan 666 | 21 | Heyu 397 | 31 | Taipingyang 891 |
| 2 | Guidan 165 | 12 | Guidan 668 | 22 | Jiafu 399 | 32 | Tiangui 88 |
| 3 | Guidan 166 | 13 | Guidan 670 | 23 | Kangnong 20 | 33 | Wanchuan 1306 |
| 4 | Guidan 203 | 14 | Guidan 673 | 24 | Longrui 888 | 34 | Wanchuan 973 |
| 5 | Guidan 550 | 15 | Guidan 902 | 25 | Longrui 999 | 35 | Yindieyu 9 |
| 6 | Guidan 551 | 16 | Guidan 903 | 26 | Lvhai 151 | 36 | Yindieyu 11 |
| 7 | Guidan 552 | 17 | Guidan 0810 | 27 | Lvhai 200 | 37 | Zhaofeng 505 |
| 8 | Guidan 553 | 18 | Guidan 6212 | 28 | Qingqing 515 | 38 | Zhaofeng 588 |
| 9 | Guidan 658 | 19 | Guiya 558 | 29 | Ruifeng 319 | 39 | Zhaoyu 200 |
| 10 | Guidan 660 | 20 | Guizhao 556 | 30 | Ruiyu 158 | 40 | Zhengda 619 |

### Morphological index

Three plants of equal growth status per pot were measured to investigate plant height and leaf area/plant (LA) by a straightedge and stem diameter through a digital vernier caliper. The LA was calculated as the followed formula:

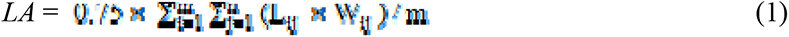

where, 0.75 was the empirical correction coefficient for LA of maize; m was the number of plants measured which was three in the study; n was the total number of leaves in the ith plant; L_ij_ and W_ij_ were the maximum leaf length and maximum leaf width of jth leaf in the ith plant.

### Preparation of enzyme extracts and determination of antioxidant enzymes

The 0.2 g fresh leaf with midrib removed in a 2.0 mL tube was grated using the high-throughput cold grinding machine (Xinyi-48N, Ningbo Xinyi Ultrasonic Equipment Co., LTD, Zhejiang, China) that was set with the speed of 55 Hz for 50 minutes under −4°C. The homogenate was blended with 2.0 mL 50 mM phosphate buffer solution (PBS, pH 7.8) that contained 1.0% polyvinyl pyrrolidone and oscillated fully. Subsequently, the homogenate was centrifuged at 12,000 g for 15 min under 4°C, and the supernatant stored at 4°C ice was used for the assay of SOD, POD, CAT, APX, soluble protein and superoxide anion (O_2_^−^).

Superoxide dismutase (SOD) activity was measured based on the method of Giannopolitis and Ries (1977). The reaction solution (the reaction mixture consisted of 50 mM phosphate buffer (pH 7.8), 130 mM methionine, 750 µM NBT, 100 µM EDTA-Na_2_, 20 µM riboflavin, and distilled water) mixed with 0.1 mL enzyme solution, as well as 0.1 mL distilled water (control). The mixture solution of treatments and three controls were then placed in the light incubator for 15 min under 4000 Lux light at 30°C, and another three control were placed in the dark in the same environment. The absorbance of 50.0% photochemical reduction of NBT was measured at 560 nm using a luminometer (SpectraMax Plus^384^, Molecular Devices, CA, USA), and then the SOD activity was calculated.

The activity of POD was determined based on the slightly modified procedure of guaiacol reduction (Cakmak and Marschner, 1992). The reaction mixture was made up of 50 mL PBS (0.2 mM, pH 6.0), 28 µL guaiacol and 19 μL 30.0% hydrogen peroxide (H_2_O_2_). When the 50 μL enzyme solution was added to a 3 mL reaction mixture, the absorbance of the mixture was measured quickly at 470 nm every 30 s in 2 min by a spectrophotometer (SP-1920, Shanghai Spectral Instrument Co., LTD, China). Enzyme solution replaced with PBS as control was also assayed using the same method, and then the POD activity was calculated.

The CAT activity using ultra-violet absorption spectrometry (Luna et al., 2005) was measured based on the absorbance of the reaction mixture (100 mL 0.15 M PBS with pH 7.0) mixed with 0.1 mL enzyme solution at 240 nm every 30 s in 2 min by a spectrophotometer (SP-1920, Shanghai Spectral Instrument Co., LTD, China).

The APX activity was measured using the method of ASA (ascorbic acid) oxidation (Nakano and Asada, 1987). The 2.6 mL PBS (50 mM, pH 7.0) containing 0.1 mM EDTA-Na_2_, 0.15 mL 5 mM ascorbic acid, and 0.15 mL 20 mM H_2_O_2_ were successively added into 0.1 mL enzyme solution, and then the mixture was measured quickly at 290 nm every 30 s in 2 min by a spectrophotometer (SP-1920, Shanghai Spectral Instrument Co., LTD, China). Pre-cooled PBS was used instead of enzyme liquid as control.

### Measurement of the reactive oxygen species

The method of hydroxylamine oxidation was adopted to estimate the generation rate of superoxide anion O_2_^−^ (Lukatkin, 2002). The 0.5 mL PBS (0.05M, pH7.8) and 1.0 mL hydroxylamine hydrochloride (10 mM) were added to 0.5 mL supernatant same as that of SOD. Thereafter, the reaction was incubated at 25 °C for 1 h. Subsequently, 1.0 mL para-aminobenzoic acid (17 mM) and 1.0 mL α-naphthylamine (7 mM) were added in order and then incubated at 25 °C for 20 min. The absorbance of the mixture was measured at 540 nm using a luminometer (SpectraMax Plus^384^, Molecular Devices, CA, USA), and then the rate of O_2_^−^ generation was calculated.

The H_2_O_2_ content was measured based on the slightly modified method of Velikova et al. (2000). The 0.2 g fresh leaf with midrib removed in a 2.0 mL tube was grated using the high-throughput cold grinding machine (Xinyi-48N, Ningbo Xinyi Ultrasonic Equipment Co., LTD, Zhejiang, China) that was set with the speed of 55 Hz for 50 minutes under −4°C. The homogenate was blended with 2.0 mL trichloroacetic acid (0.1%, W/V) and centrifuged at 12,000 g for 15 min under 4°C. The 0.5 mL supernatant mixed with 10 mM PBS (pH7.0) and 1.0 M potassium iodide. The mixture was assayed at 390 nm using a luminometer (SpectraMax Plus^384^, Molecular Devices, CA, USA), and then H_2_O_2_ content was calculated based on the standard curve of H_2_O_2_.

### Assessment of the osmotic adjustment substances

The soluble protein content was measured according to the procedure of coomassie brilliant blue G-250 staining (Bradford, 1976). 0.1 mL supernatant that was extracted as same as that of SOD mixed with 5.0 mL coomassie brilliant blue G-250 has measured the absorbance of 595 nm using a luminometer (SpectraMax Plus^384^, Molecular Devices, CA, USA).

Microcalorimetry was used to extract soluble sugar (Leyva et al., 2008). 0.2 g fresh leaf grated was incubated with 1.6 mL distilled water and then boiled for 40 min. Thereafter, the liquid was decolored by activated carbon. The supernatant was got after centrifuging at 10,000 g for 5 min under 4°C. The 50.0 μL supernatant mixed with 50.0 μL ethyl alcohol and 0.15 mL anthrone reagent was boiled for 10 min. The absorbance of 620 nm for soluble sugar was measured using a luminometer (SpectraMax Plus384, Molecular Devices, CA, USA).

The proline content was assayed by the procedure of (Bates et al., 1973). The grated fresh leaf (0.2g) was homogenized in 2.0 mL sulfosalicylic acid (Xinyi-48N, Ningbo Xinyi Ultrasonic Equipment Co., LTD, Zhejiang, China) and boiled for 10 min. The homogenate was centrifuged at 10,000 g for 10 min under 4°C to extract supernatant for further analysis. 0.2 mL glacial acetic acid, 0.4 mL acid ninhydrin and 0.2 mL sulfosalicylic acid were added to 0.2 mL supernatant and then boiled for 30 min. 0.4 mL toluene was added to the mixture. After shocking and standing, the supernatant was centrifuged at 900 g for 5 min under 4°C and then measured by a luminometer (SpectraMax Plus384, Molecular Devices, CA, USA).

### Resistance analysis

The resistance coefficient for drought and flooding was the ratio of treatment value to CK value (Sun, et al., 2021). The membership function value μ(X_ij_) and D value for each maize variety was calculated by the formulas proposed by Zou et al. (2020) as follows:

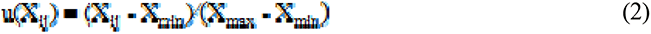

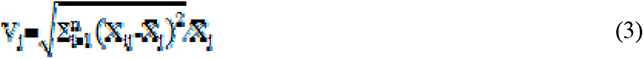

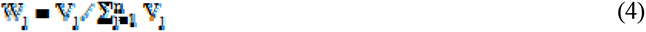

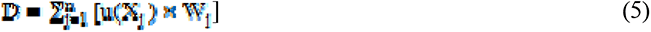

Where X_ij_ was the value of ith variety in the jth indicator; X_min_, X_max_, V_j_ and 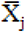 indicated the minimum value, maximum value, standard deviation coefficient and average value for the jth indicator of all the varieties, respectively; and W_j_ represented the weight of the jth indicator in all indicators.

### Data analysis

The two-factors analysis of variance (ANOVA) was analyzed by SAS 9.2 (Sas Institute, 2009), and the significant difference between treatments was identified based on the LSD test at *p* ≤0.05. The software of SPSS statistics v. 21 (IBM Inc., Armonk, N.Y., USA) performed the PCA and stepwise regression analysis. Meanwhile, OriginPro 2021 (OriginLab Inc., Northampton, Massachusetts., USA) was used to draw all figures.

## Results

### Maize-seedlings growth trait indices in response to water stress

The statistical analysis for osmotic adjustment substances, antioxidative enzymes and morphological characteristics of 40 varieties was shown in Table 2. No matter what kind of water stress was applied, all indexes were significantly affected by water stress, and a large variation of different indexes was found in different varieties of maize (*p* < 0.01, Table 2 and Figure S1-S7). The coefficient of variation in all indexes ranged from 7.96% to 58.33%. Meanwhile, a wide range of coefficient of variation from 7.99% to 141.67% was identified in the resistance coefficient, in which the coefficient of variation in osmotic adjustment substances under drought stress was higher than that of flooding stress while other indicators existed an opposite trend. For a comparable mean in all indexes, there was a significant increment in osmotic adjustment substances and antioxidative enzymes (except APX) for maize in drought stress compared with those in CK and flooding stress (Table 2). However, not all indicators in all varieties followed the above same trend, for example, soluble sugar and CAT severally in 30.0% and 22.5% of varieties under drought stress were decreased compared with CK (Figure S2, S3, S6). Correspondingly, under flooding stress, only soluble sugar, POD, and APX in flooding stress were higher than those in CK, and over 50.0% of varieties in other indicators especially in proline and SOD had lower contents than CK (Figure S3 and S4). Significant reductions in LA, plant height and stem diameter were detected in either drought or flooding stress compared with those in CK (*p* < 0.05), in which flooding stress contributed more reductions in LA and plant height compared with drought stress (Table 2 and S1). Whereas, when subjected to flooding stress, about 55.0% of varieties maintained a larger stem diameter compared with drought stress (Table S1). As a consequence, the average drought-resistance coefficients in osmotic adjustment substances and antioxidative enzymes were over 1.00 while flooding-resistance coefficients in those indexes except soluble sugar, POD and APX were less than 1.00. Whether drought- or flooding-resistance coefficients in morphological characteristics were all lower than 0.9 (Table 2).

**Table 2.** Statistical analysis of main indexes in maize under water stress.

| Index | Water stress | Variation range | Mean ± SD | CV ( % ) | $R^2$ | RC mean | CV of RC ( % ) |
| --- | --- | --- | --- | --- | --- | --- | --- |
| Protein (mg/g FW) | Well water | 1.33-2.72 | 2.00±0.37b | 18.56 | 0.96** | - | - |
| | Drought | 1.45-3.31 | 2.47±0.46a | 18.54 | | 1.26 | 18.32 |
| | Flooding | 1.26-2.65 | 1.89±0.30c | 16.04 | | 0.98 | 24.26 |
| Soluble sugar (mg/g FW) | Well water | 3.24-13.28 | 7.14±2.20c | 30.83 | 0.99** | - | - |
| | Drought | 5.03-20.57 | 11.77±4.76a | 40.45 | | 1.84 | 61.58 |
| | Flooding | 5.65-22.70 | 10.47±4.05b | 38.70 | | 1.59 | 45.99 |
| Proline (μmol/g FW) | Well water | 12.67-30.53 | 18.23±4.57b | 25.04 | 0.99** | - | - |
| | Drought | 11.11-86.71 | 25.48±14.86a | 58.33 | | 1.46 | 59.89 |
| | Flooding | 6.90-19.27 | 13.36±3.11c | 23.31 | | 0.77 | 30.63 |
| SOD (U/g FW/min) | Well water | 11.13-19.94 | 16.23±1.88b | 11.57 | 0.99** | - | - |
| | Drought | 10.12-20.16 | 16.78±2.32a | 13.84 | | 1.05 | 18.28 |
| | Flooding | 8.22-15.46 | 12.58±1.90c | 15.08 | | 0.78 | 14.39 |
| POD (U/mg FW/min) | Well water | 22.95-52.29 | 36.29±6.77c | 18.64 | 0.96** | - | - |
| | Drought | 28.14-65.45 | 40.27±7.85a | 19.48 | | 1.12 | 16.67 |
| | Flooding | 27.59-60.78 | 37.21±6.78b | 18.23 | | 1.05 | 20.07 |
| CAT (U/mg FW/min) | Well water | 0.25-2.28 | 1.05±0.51b | 48.52 | 0.98** | - | - |
| | Drought | 0.31-2.62 | 1.08±0.44a | 40.62 | | 1.24 | 55.62 |
| | Flooding | 0.23-1.91 | 0.84±0.45c | 53.41 | | 1.08 | 81.51 |
| APX (mM/g FW/min) | Well water | 0.37-6.73 | 2.83±1.57b | 55.57 | 0.99** | - | - |
| | Drought | 0.44-6.85 | 3.98±1.70a | 42.68 | | 2.29 | 108.61 |
| | Flooding | 0.54-7.23 | 4.02±1.69a | 41.98 | | 2.57 | 141.67 |
| Leaf area/plant (cm<sup>2</sup>) | Well water | 93.41-226.94 | 154.45±38.20a | 24.74 | 0.96** | - | - |
| | Drought | 80.48-162.62 | 110.33±19.68b | 17.84 | | 0.75 | 23.98 |
| | Flooding | 66.34-157.09 | 108.55±21.37c | 19.69 | | 0.73 | 21.84 |
| Plant height (cm) | Well water | 51.81-76.27 | 64.08±6.14a | 9.58 | 0.95** | - | - |
| | Drought | 45.45-67.13 | 53.52±4.52b | 8.44 | | 0.84 | 10.35 |
| | Flooding | 41.10-62.40 | 52.26±4.96c | 9.48 | | 0.82 | 12.08 |
| Stem diameter (mm) | Well water | 4.60-6.78 | 5.56±0.56a | 10.13 | 0.94** | - | - |
| | Drought | 3.70-5.45 | 4.79±0.38b | 7.96 | | 0.87 | 7.99 |
| | Flooding | 3.53-6.10 | 4.99±0.58b | 11.58 | | 0.90 | 10.79 |

CV, Coefficient of variation; SD, Standard deviation; RC, Resistance coefficient; \*\*, Significant difference at $p < 0.01$ ; Means in a indicator with the same letter are not significantly different at $p > 0.05$ .

### The accumulation of reactive oxygen species under water stress

Reactive oxygen species including H_2_O_2_ and O_2_^−^ were quantified and compared with regard water stress. As a consequence, a bulk of ROS was accumulated in water stress compared with CK, which was affected significantly not only by water stress but also by variety (*p* < 0.01, Figure 2, Figure S8 and S9). The ROS increased by 106.90% in flooding while only by 29.45% in drought compared with CK, of which about 29.23% and 107.12% of the increment rate were induced in H_2_O_2_ and O_2_^−^, respectively (Figure 2). The average H_2_O_2_ and O_2_^−^ contents were 0.36 μmol/g FW and 31.57μg/g FW in drought and 0.36 μmol/g FW and 67.64 μg/g FW in flooding, respectively. At the same time, the contents of H_2_O_2_ and O_2_^−^ among most varieties under drought stress were significantly higher than those under flooding stress (*p* < 0.015). The highest and lowest ROS were produced in Guidan 553 and Qingqing 515 under drought stress, as well as Lvhai 200 and Yindieyu 11 under flooding stress, respectively. The variety of Guidan 0810 under both drought and flooding stresses always kept the lowest increment rate in ROS.

**Figure 2.**
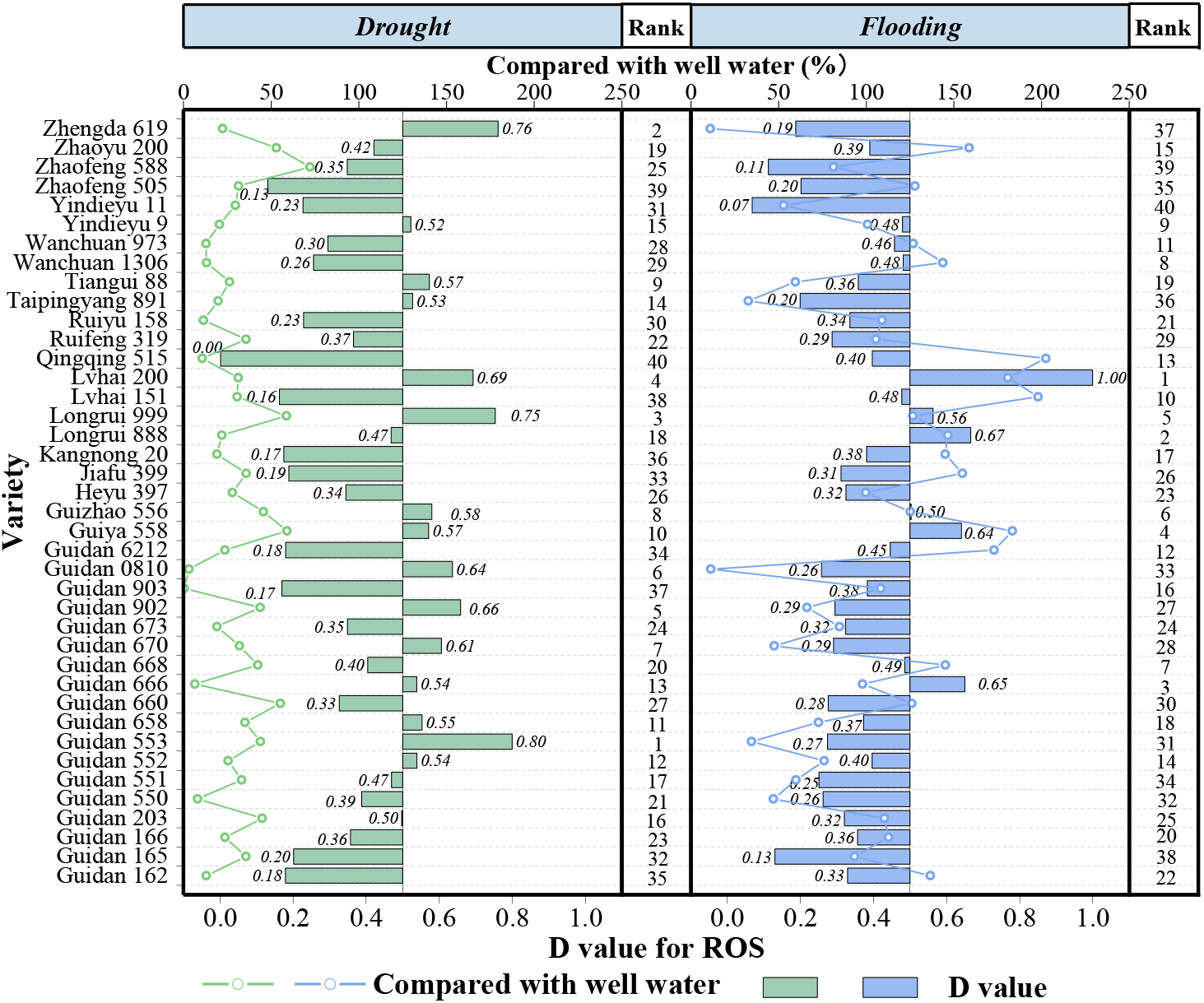
The comprehensive evaluation (D) value for reactive oxygen species (ROS) and the increment of different maize varieties under water stress compared with well water. The numbers represent the averages.

### Water tolerance is evaluated by comprehensive evaluation (D) value and resistance coefficient

A single physiological index could not be used as an indicator of water resistance in crops, but the D value played a particularly important role in evaluating the water resistance of crops. The D value was calculated based on all the target traits in the study as shown in Table 3, and then could be utilized to evolve strategies to assess water stress (Table 3). A higher D value corresponded to a higher value in all traits and thus corresponded to higher water resistance. The D value was significantly positively correlated with the resistance coefficient (*R*^*2*^ = 0.33, *p* = 0.003), and a significant positive correlation between the rank of both were also found (*R*^*2*^ = 0.58,*p* < 0.001). However, the D value and resistance coefficient were not exactly in the same rank. When maize was subjected to drought stress, the varieties of Guidan 670, Guidan 902, Guidan 0810, Guidan 673, and Guidan 668 had higher D values over 0.54; others’ D values were lower than 0.54. Meanwhile, the lower D values in Zhaofeng 505, Tiangui 88, Guidan 903, Qingqing 515, Lvhai 200, Lvhai 151, Taipingyang 891, Wanchuan 973, Longrui 888, and Heyu 397 increased successively, but their D values were all less than equal to 0.30. The drought-resistance coefficient in a variety with a high D value was also high, and the variety with a low D value also had a low drought-resistance coefficient. When maize was subjected to flooding stress, the D value of Guidan 551, Taipingyang 891, Guidan 668, Wanchuan 973, and Guidan 6212 decreased successively and were all greater than the others’, but only Taipingyang 891, Guidan 668 and Wanchuan 973 had the higher flooding-resistance coefficient. The D value of Lvhai 151 and Zhaofeng 505 were all 0.22 which were significantly lower than others under flooding stress, of which Zhaofeng 505 had the weakest flooding resistance in those varieties with a 0.89 flooding-resistance coefficient (*p* < 0.05). The drought- and flooding-resistance coefficients of 80.0% and 57.5% of varieties were greater than 1.0, respectively, and 75.0% of varieties had higher drought-resistance coefficients compared with their flooding-resistance coefficient.

**Table 3.** Comprehensive evaluation (D) value and resistance coefficient of maize varieties under water stress.

| Variety | Drought |  |  |  | Flooding |  |  |  | Variety | Drought |  |  |  | Flooding |  |  |  |
| --- | --- | --- | --- | --- | --- | --- | --- | --- | --- | --- | --- | --- | --- | --- | --- | --- | --- |
|  | D value | Rank | DRC | Rank | D value | Rank | FRC | Rank |  | D value | Rank | DRC | Rank | D value | Rank | FRC | Rank |
| Guidan 162 | 0.42 | 13 | 2.21 | 1 | 0.54 | 6 | 2.42 | 2 | Heyu 397 | 0.30 | 31 | 0.91 | 37 | 0.33 | 33 | 0.78 | 39 |
| Guidan 165 | 0.42 | 12 | 1.43 | 11 | 0.40 | 26 | 1.08 | 16 | Jiafu 399 | 0.37 | 21 | 1.01 | 32 | 0.37 | 32 | 0.83 | 36 |
| Guidan 166 | 0.50 | 8 | 1.63 | 7 | 0.40 | 27 | 1.16 | 9 | Kangnong 20 | 0.34 | 25 | 1.97 | 3 | 0.43 | 20 | 2.17 | 3 |
| Guidan 203 | 0.33 | 26 | 1.95 | 4 | 0.49 | 10 | 2.91 | 1 | Longrui 888 | 0.29 | 32 | 1.06 | 27 | 0.38 | 31 | 0.94 | 27 |
| Guidan 550 | 0.53 | 6 | 1.29 | 13 | 0.42 | 21 | 0.92 | 29 | Longrui 999 | 0.39 | 18 | 1.23 | 15 | 0.48 | 11 | 1.03 | 20 |
| Guidan 551 | 0.50 | 9 | 1.15 | 21 | 0.66 | 1 | 1.08 | 15 | Lvhai 151 | 0.28 | 35 | 2.15 | 2 | 0.22 | 40 | 1.32 | 5 |
| Guidan 552 | 0.44 | 11 | 1.23 | 14 | 0.52 | 8 | 1.07 | 17 | Lvhai 200 | 0.28 | 36 | 0.99 | 33 | 0.46 | 13 | 0.96 | 26 |
| Guidan 553 | 0.51 | 7 | 1.18 | 20 | 0.43 | 17 | 0.87 | 34 | Qingqing 515 | 0.25 | 37 | 0.96 | 35 | 0.41 | 23 | 0.94 | 28 |
| Guidan 658 | 0.40 | 17 | 1.14 | 22 | 0.43 | 19 | 0.99 | 24 | Ruifeng 319 | 0.42 | 14 | 1.11 | 24 | 0.29 | 38 | 0.87 | 33 |
| Guidan 660 | 0.41 | 15 | 1.21 | 18 | 0.31 | 36 | 0.83 | 37 | Ruiyu 158 | 0.33 | 27 | 1.14 | 23 | 0.41 | 24 | 1.01 | 22 |
| Guidan 666 | 0.33 | 28 | 1.03 | 29 | 0.52 | 7 | 1.14 | 10 | Taipingyang 891 | 0.28 | 34 | 0.95 | 36 | 0.64 | 2 | 1.23 | 6 |
| Guidan 668 | 0.54 | 5 | 1.54 | 9 | 0.62 | 3 | 1.51 | 4 | Tiangui 88 | 0.23 | 39 | 0.87 | 40 | 0.45 | 14 | 0.99 | 25 |
| Guidan 670 | 0.75 | 1 | 1.51 | 10 | 0.47 | 12 | 0.80 | 38 | Wanchuan 1306 | 0.35 | 22 | 1.22 | 17 | 0.40 | 25 | 1.09 | 14 |
| Guidan 673 | 0.59 | 4 | 1.66 | 6 | 0.44 | 15 | 1.06 | 18 | Wanchuan 973 | 0.29 | 33 | 0.89 | 38 | 0.61 | 4 | 1.18 | 8 |
| Guidan 902 | 0.62 | 2 | 1.54 | 8 | 0.39 | 29 | 1.09 | 13 | Yindieyu 9 | 0.35 | 23 | 1.02 | 30 | 0.39 | 28 | 0.90 | 30 |
| Guidan 903 | 0.25 | 38 | 1.06 | 28 | 0.41 | 22 | 1.02 | 21 | Yindieyu 11 | 0.31 | 29 | 1.02 | 31 | 0.43 | 18 | 1.06 | 19 |
| Guidan 0810 | 0.62 | 3 | 1.83 | 5 | 0.43 | 16 | 1.10 | 12 | Zhaofeng 505 | 0.18 | 40 | 0.88 | 39 | 0.22 | 39 | 0.89 | 31 |
| Guidan 6212 | 0.49 | 10 | 1.20 | 19 | 0.57 | 5 | 1.01 | 23 | Zhaofeng 588 | 0.34 | 24 | 1.30 | 12 | 0.38 | 30 | 1.19 | 7 |
| Guiya 558 | 0.38 | 20 | 0.97 | 34 | 0.30 | 37 | 0.71 | 40 | Zhaoyu 200 | 0.40 | 16 | 1.23 | 16 | 0.31 | 35 | 0.86 | 35 |
| Guizhao 556 | 0.31 | 30 | 1.09 | 26 | 0.51 | 9 | 1.14 | 11 | Zhengda 619 | 0.38 | 19 | 1.11 | 25 | 0.32 | 34 | 0.88 | 32 |

DRC, Drought-resistance coefficient; FRC, Flooding-resistance coefficient.

### Key drought or flooding tolerance indicators identified through principal component analysis

Through PCA, the standardized data tested by KMO was 0.64 (> 0.6) with *p* = 0 (Bartlett. Sphericity test), so there was a correlation among the indicators. Four top principal components were extracted based on the eigenvalue over 1.0, marked as F_1_ to F_4_, which explained separately 25.45%, 18.72%, 11.38%, and 11.25% of the total variation and together represented 66.79% of the information of the 10 indicators (Table 4, Figure S10). The F_1_ was related to proline, SOD, POD, and CAT which could be defined as an antioxidant factor. The F_2_ was related to morphological indexes such as LA, plant height, and stem diameter, which could be defined as a comprehensive morphological factor. The maximum loading in F_3_ was found in soluble sugar and protein that could be defined as an osmotic adjustment factor. The F_4_ was mainly related to APX, indicating that it played a great role in antioxidant indicators under water stress. Based on the weight of principal component, the equations for four and total principal component (F) were in order as followed: *F*_*1*_ = 0.275*X*_*1*_ + 0.235*X*_*2*_ + 0.502*X*_*3*_ + 0.483*X*_*4*_ + 0.393*X*_*5*_ + 0.324*X*_*6*_ + 0.114*X*_*7*_ + 0.029*X*_*8*_ + 0.076*X*_*9*_ − 0.182*X*_*10*_ (6), *F*_*2*_ = 0.069*X*_*1*_ + 0.010*X*_*2*_ − 0.057*X*_*3*_ + 0.034*X*_*4*_ − 0.081*X*_*5*_ + 0.133*X*_*6*_ + 0.060*X*_*7*_ + 0.607*X*_*8*_ + 0.586*X*_*9*_ + 0.501*X*_*10*_ (7), *F*_*3*_ = −0.731*X*_*1*_ + 0.557*X*_*2*_ + 0.1441*X*_*3*_ − 0.103*X*_*4*_ + 0.135*X*_*5*_ + 0.052*X*_*6*_ + 0.234*X*_*7*_ + 0.172*X*_*8*_ −0.008*X*_*9*_ − 0.136*X*_*10*_ (8), *F*_*4*_ = 0.007*X*_*1*_ + 0.195*X*_*2*_ + 0.150*X*_*3*_ + 0.174*X*_*4*_ + 0.089*X*_*5*_ + 0.237*X*_*6*_ + 0.234*X*_*7*_ + 0.200*X*_*8*_ + 0.172*X*_*9*_ + 0.040*X*_*10*_ (10), *F* = 0.381*F*_1_ + 0.280*F*_2_ + 0.170*F*_3_ + 0.168*F*_4_ (11). Where, *X*_*1*_, *X*_*2*_, *X*_*3*_, *X*_*4*_, *X*_*5*_, *X*_*6*_, *X*_*7*_, *X*_*8*_, *X*_*9*_, and *X*_*10*_ represented 10 indicators, namely, soluble protein, sugar, proline, SOD, POD, CAT, APX, LA, plant height, and stem diameter. Based on the comprehensive score of F, there were nine varieties with a higher score over 1.0 such as Guidan 902, Guidan 668, Guidan 673, Guidan 6212, Guidan 0810 while lower scores less than −0.5 were found in Zhaofeng 505, Tiangui 88, Guidan 903 and Taipingyamg 891 under drought stress (Figure 3). Meanwhile, under flooding stress, the higher comprehensive scores were detected in the varieties of Guidan 668, Wanchuan 973, Guidan 551, and Taipingyamg 891, and lower scores less than −1.0 were in Zhaofeng 505, Lvhai 151, Guidan 660, Heyu397 and Guiya 558 (Figure 4). The F_1_ played a dominant role in the comprehensive score (contribution rate 25.45%). The comprehensive score of 82.5% varieties in drought stress was higher than that in flooding stress. In addition, 55.0% of maize varieties in drought stress had a comprehensive score of F higher than zero, but only 27.5% of maize varieties in flooding stress.

**Table 4.** Loading matrix and the variance contribution rate of the principal component under water stress.

| Index | Principal component |  |  |  | Weight of principal component |  |  |  | Total |
| --- | --- | --- | --- | --- | --- | --- | --- | --- | --- |
|  | F<sub>1</sub> | F<sub>2</sub> | F<sub>3</sub> | F<sub>4</sub> | F<sub>1</sub> | F<sub>2</sub> | F<sub>3</sub> | F<sub>4</sub> | F |
| Protein (X<sub>1</sub>) | 0.438 | 0.095 | -0.78 | 0.045 | 0.275 | 0.069 | -0.731 | 0.042 | 0.007 |
| sugar (X<sub>2</sub>) | 0.567 | 0.013 | 0.594 | -0.238 | 0.355 | 0.010 | 0.557 | -0.224 | 0.195 |
| Proline (X<sub>3</sub>) | 0.801 | -0.078 | -0.154 | -0.002 | 0.502 | -0.057 | -0.144 | -0.002 | 0.150 |
| SOD (X<sub>4</sub>) | 0.771 | 0.047 | -0.11 | -0.011 | 0.483 | 0.034 | -0.103 | -0.010 | 0.174 |
| POD (X<sub>5</sub>) | 0.627 | -0.111 | 0.144 | -0.387 | 0.393 | -0.081 | 0.135 | -0.365 | 0.089 |
| CAT (X<sub>6</sub>) | 0.517 | 0.182 | 0.055 | 0.423 | 0.324 | 0.133 | 0.052 | 0.399 | 0.237 |
| APX (X<sub>7</sub>) | 0.182 | 0.082 | 0.25 | 0.845 | 0.114 | 0.060 | 0.234 | 0.797 | 0.234 |
| LA (X<sub>8</sub>) | 0.046 | 0.831 | 0.183 | -0.066 | 0.029 | 0.607 | 0.172 | -0.062 | 0.200 |
| Plant height (X<sub>9</sub>) | 0.122 | 0.802 | -0.009 | -0.126 | 0.076 | 0.586 | -0.008 | -0.119 | 0.172 |
| Stem diameter (X<sub>10</sub>) | -0.29 | 0.685 | -0.145 | -0.049 | -0.182 | 0.501 | -0.136 | -0.046 | 0.040 |
| Eigenvalue | 2.545 | 1.872 | 1.138 | 1.125 | - | - | - | - | - |
| CR(%) | 25.448 | 18.724 | 11.375 | 11.245 | - | - | - | - | - |
| CCR (%) | 25.448 | 44.172 | 55.547 | 66.792 | - | - | - | - | - |
| Weight | 0.381 | 0.280 | 0.170 | 0.168 | - | - | - | - | - |

F<sub>1</sub>, F<sub>2</sub>, F<sub>3</sub>, F<sub>4</sub>, four top principal components; SOD, superoxide dismutase; POD, peroxidase; CAT, catalase; APX, anticyclohexate peroxidase; LA, leaf area/plant; CR, contribution rate; CCR, cumulative contribution rate

**Figure 3.**
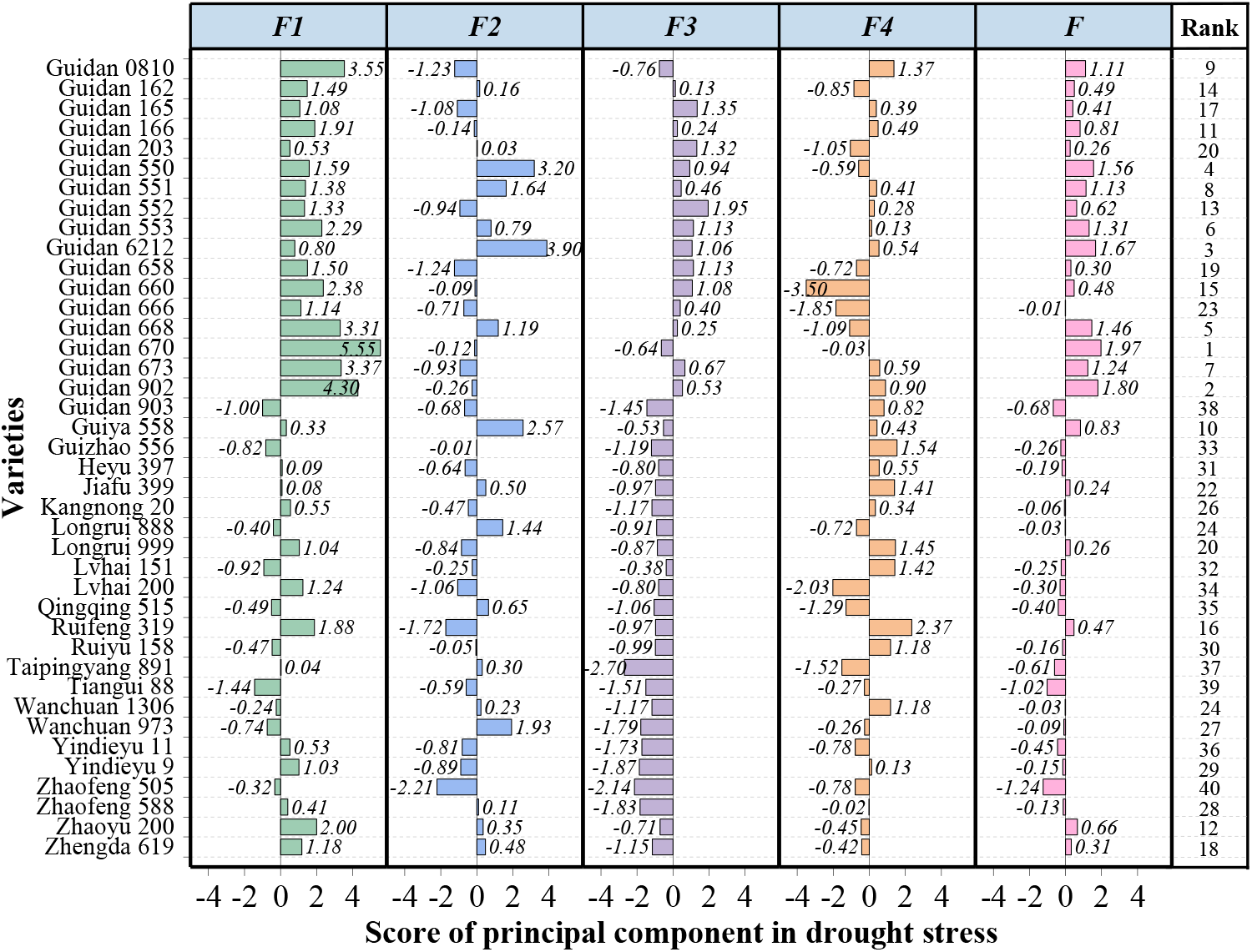
The score of principal components for all the varieties under drought stress. F_1_, F_2_, F_3_ and F_4_ are the four top principal components. The numbers represent the scores.

**Figure 4.**
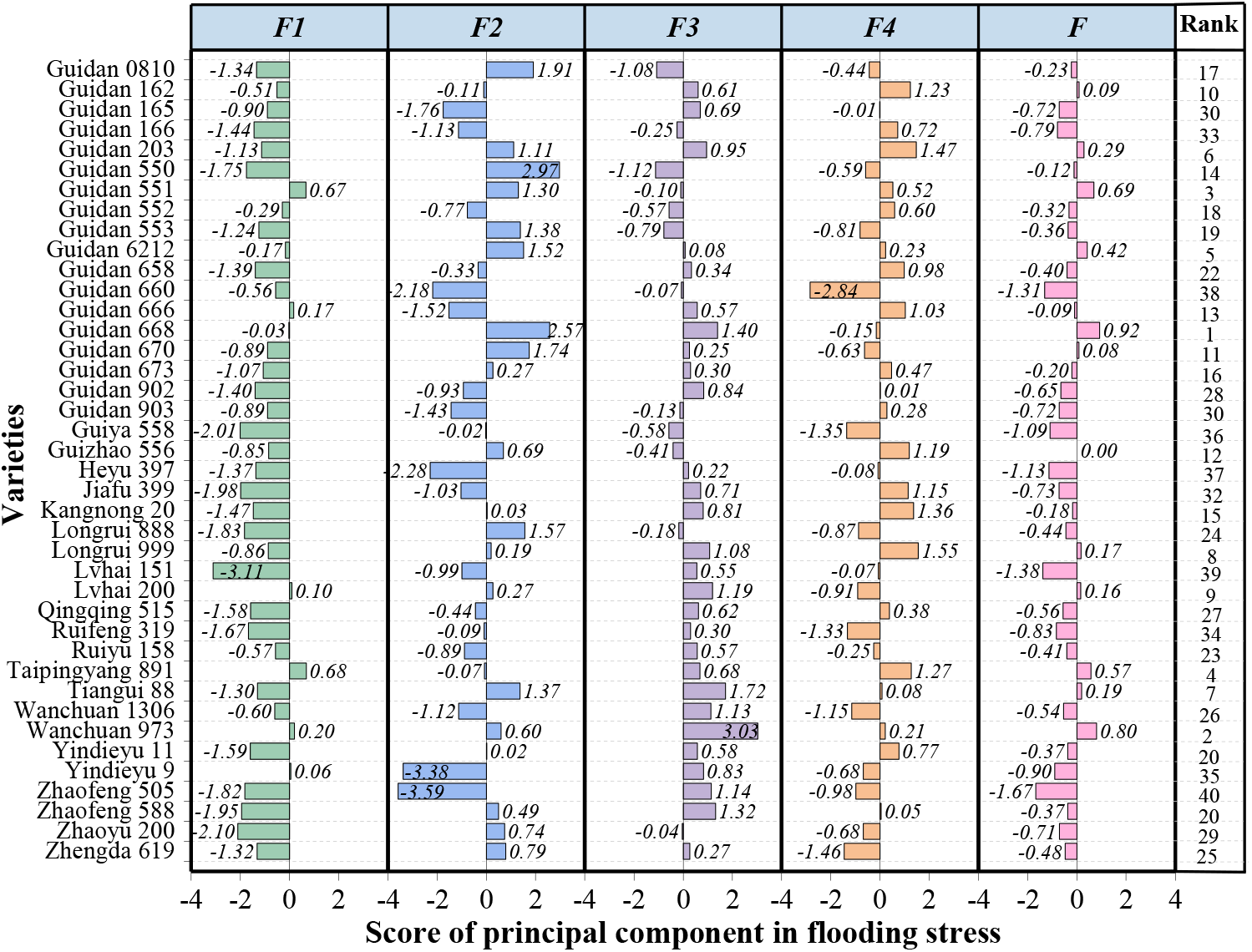
The score of principal components for all the varieties under flooding stress. F_1_, F_2_, F_3_ and F_4_ are the four top principal components. The numbers represent the scores.

**Figure 5.**
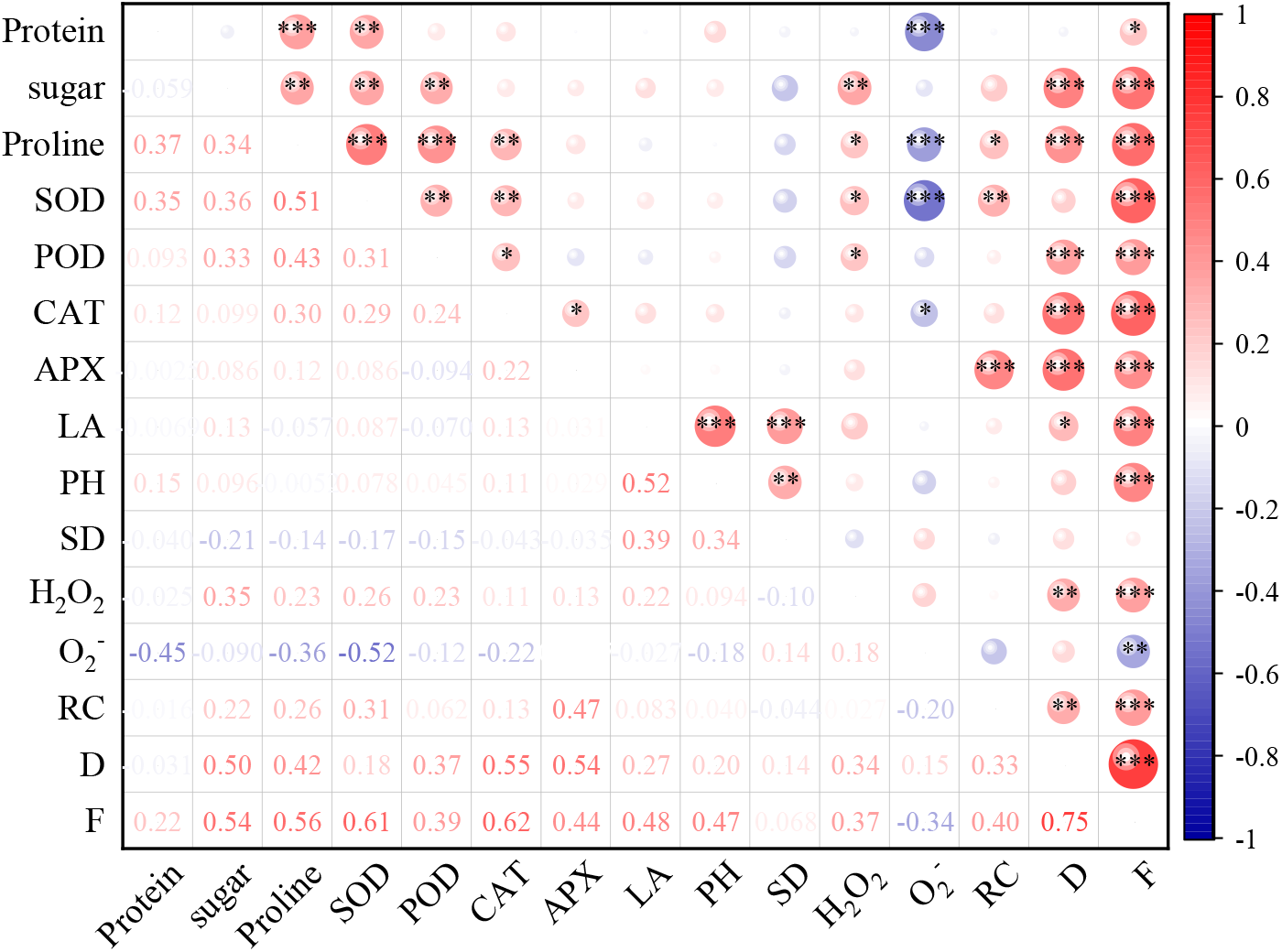
Correlation analysis between all the traits indices. *, *p* ≤ 0.05; **, *p* ≤ 0.01; ***, *p* ≤ 0.001; SOD, superoxide dismutase; POD, peroxidase; CAT, catalase; APX, anticyclohexate peroxidase; LA, leaf area/plant; RC, resistance coefficient; H_2_O_2_, hydrogen peroxide; O_2_^−^, superoxide anion; D, comprehensive evaluation value; F, comprehensive score of principal components

Appropriate indicators were selected by stepwise regression analysis to comprehensively assess the flooding and drought tolerance of the maize varieties. The 10 trait indicators and comprehensive D value were set as the independent and dependent variables, respectively. The optimum regression equation for drought stress was *D*_*1*_ = −0.458 + 0.037*X*_*1*_ + 0.001*X*_*2*_ + 0.003*X*_*3*_ + 0.005*X*_*4*_+ 0.002*X*_*5*_ + 0.066*X*_*6*_ + 0.025*X*_*7*_ + 0.001*X*_*8*_ + 0.001*X*_*9*_ + 0.016 *X*_*10*_ (12, *R*^*2*^ = 0.999, *p* < 0.001) and for flooding was *D*_*2*_ = −0.527 + 0.043*X*_*1*_ + 0.009*X*_*2*_ + 0.007*X*_*3*_ + 0.008*X*_*4*_ + 0.002*X*_*5*_ + 0.127*X*_*6*_ + 0.026*X*_*7*_ + 0.001*X*_*8*_ + 0.002*X*_*9*_ + 0.017*X*_*10*_ (13, *R*^*2*^ = 0.999, *p* < 0.001). The path coefficient of X_1_, X_2_, X_3_, X_4_, X_5_, X_6_, X_7_, X_8_, X_9_, X_10_ was 0.138, 0.381, 0.355, 0.089, 0.127, 0.237, 0.349, 0.137, 0.044, 0.047 in drought and 0.125, 0.354, 0.214, 0.148, 0.155, 0.549, 0.417, 0.178, 0.084, 0.094 in flooding, respectively. All the 10 indicators had significant effects on the comprehensive D value and could be used as key trait indices of a comprehensive evaluation for drought and flooding tolerance of maize.

### The correlation analysis between all the trait indices

Through the correlation analysis, the D value was significantly and positively correlated with other indicators except for protein, SOD, PH, stem diameter, and O_2_^−^ (Figure 4). Meanwhile, significant positive correlations were observed between the F value and all the other trait indices except stem diameter (*p* ≤ 0.05). Yet somehow, the resistance coefficient was only positively correlated with proline, SOD, APX, D, and F values. There were significant positive correlations between POD, CAT, proline, and SOD (*p* ≤ 0.05). The O_2_^−^ had significant negative effects on protein, proline, SOD, CAT, and F value (*p* ≤ 0.05).

### Evaluation of drought and flooding tolerance in maize by cluster analysis

Cluster analysis results of 40 maize varieties based on comprehensive D and F values were divided into five groups including high resistance, resistance, moderate resistance, weak sensitivity and sensitivity to drought or flooding (Figure 6). Group V including varieties of Guidan 902, Guidan 0810, Guidan 673, and Guidan 670 which had the highest D values and top nine F values were confirmed to be the high resistance to drought stress (Figure 3, Table 3). Furthermore, their drought-resistance coefficient was greater than 1.50. Besides, the D and F values in group IV comprised of Guidan 6212, Guidan 668, Guidan 550, Guidan 553, Guidan 551, and Guidan 166 also still ranked top 11, which showed drought resistance among all the varieties. Moreover, varieties of Zhaofeng 505 and Tiagui 88 in group III had the lowest tolerance and highest sensitivity to drought stress, which had the lowest D and F values. Under flooding stress, the high flooding resistance was categorized into group II involving varieties of Guidan 668, Taipingyang 891, Wanchuan 973, and Guidan 551, whose D and F values raked top four were separately greater than 0.60 and 0.55, while the flooding-resistance coefficient was more than 1.0 (Figure 4, Table 3). Variety of Guidan 668 had the highest flooding-resistance coefficient among the four varieties. Interestingly, varieties of Zhaofeng 505 in group IV also exhibited the greatest sensitivity to flooding stress (0.22 of D value, −1.67 of F value, and 0.89 of flooding-resistance coefficient).

**Figure 6.**
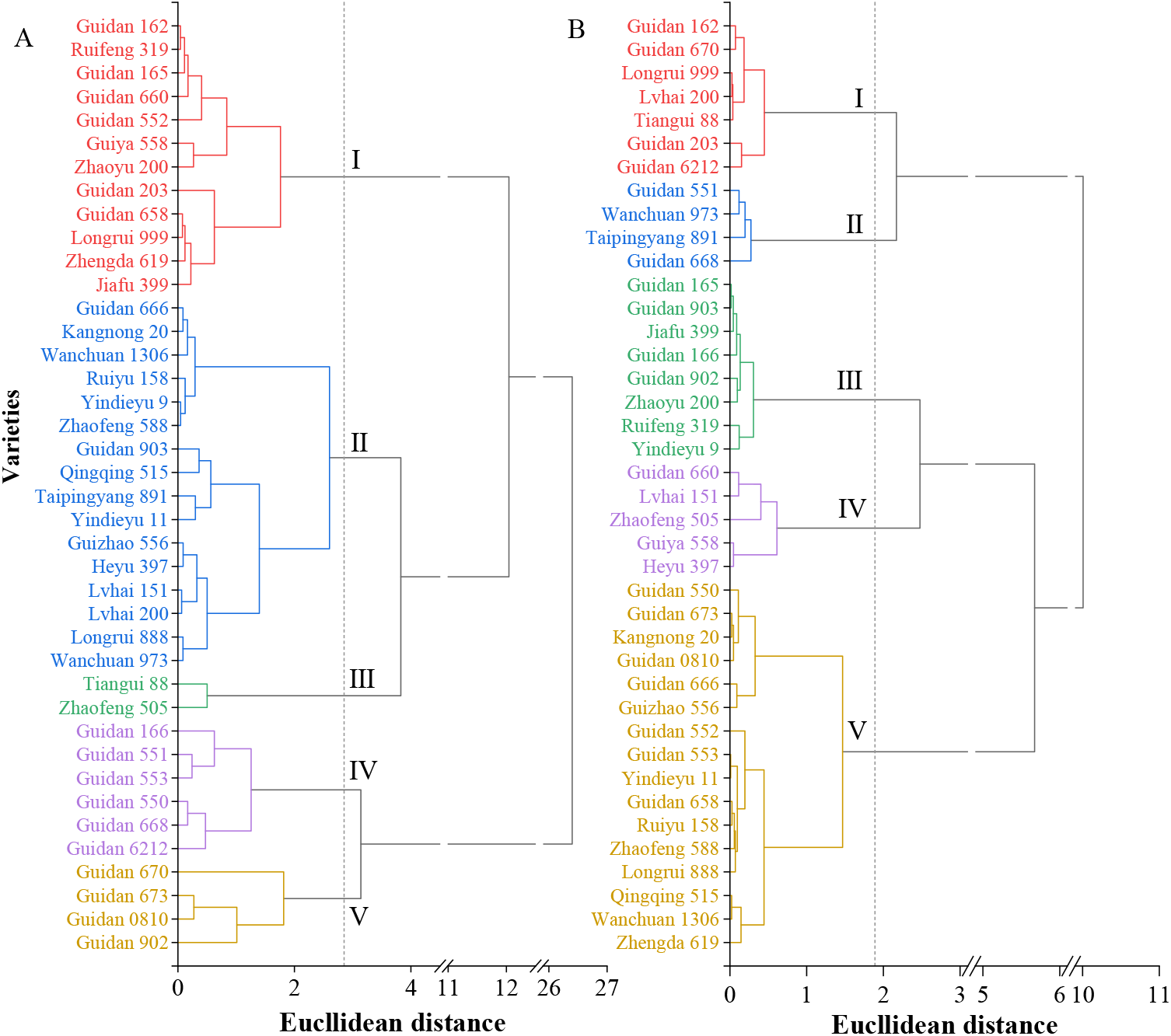
Cluster analysis for the drought (A) and flooding (B) tolerance of maize varieties based on comprehensive evaluation value and F values.

## Discussion

Maize in south China, ranking second in production acreage among crops, suffers serious damage from drought and flooding stress leading to a much lower grain production than northern area (National Bureau of Statistics of China 2010-2018). Screening and identification of drought or flooding-tolerant genotypes is the core task for breeding maize. It is one of the main objectives of drought or flooding-tolerant breeding to screen simple and effective identification indexes. A series of morphology, physiology, and biochemistry under water stress are changed, which can be applied in the resistance evaluation of germplasm (Chen et al., 2012 and 2016; Anjum et al., 2016; Wijewardana et al., 2016). Plant height, transpiration rate, and chlorophyll were detected as the key representative indicators to evaluate the drought resistance of cotton (Sun et al., 2021). Previous studies have identified nine typical indicators including proline, SOD, MDA, CAT, and so on from 18 physiological indices to screen the drought resistance variety of cotton (Zou et al., 2020). Additionally, the POD activity could also be used as a typical indicator for screening waterlogging tolerance at the maize seedling stage (LIU et al., 2010). Those scholars have developed different representative indicators from one or two aspects of morphology, physiology, and biochemistry for evaluating water stress resistance for different crops, but now only partial indicators are adaptability, and few unified and systematic indicators are developed to evaluate water stress resistance, particularly in flooding tolerance of maize as more researches focused on its drought resistance. In our study, morphological indicators (plant height, LA, and stem diameter) combined with biochemical indexes (ROS, SOD, POD, APX, CAT, proline, soluble sugar, and protein), in a total of 10 indicators, were adapted to evaluate drought and flooding resistance at the third-leaf stage of maize. Maize was susceptible to water stress, and its morphological indicators showed a considerable reduction when it suffered from drought or flooding stress. Moreover, osmotic adjustment and antioxidant enzyme activities play an essential role in resisting abiotic stress (Egert and Tevini, 2002; Fang and Xiong, 2015; Javadi et al., 2017). We found that all indicators performed significant differences in different water stress and varieties categories based on the variance analysis. Simultaneously, most coefficients of variation under the control were lower than that of the drought stress, while higher than that of flooding stress, demonstrating that the maize cultivars types selected in the study were abundant, and the results are quite representative.

It is a novel concept that most maize cultivars have different susceptibility in response to drought and flooding stress and have stronger drought resistance as opposed to flood resistance because the evaluation for drought and flooding resistance of maize germplasm had not been comparatively researched previously. As a consequence, drought stress results in a more pronounced increment in osmotic adjustment substances and antioxidative enzymes than well-watered treatment (Nikolaeva et al., 2015; Aghaie et al., 2018). In contrast, most indicators among the varieties such as protein, proline, SOD, and CAT were attenuated under flooding stress. Correspondingly, the drought-resistance coefficient in most varieties was higher than 1.0, while the flooding-resistance coefficient was less than 1.0. On the other hand, flooding stress also can lead to the increase of soluble sugar, POD, and APX among most cultivars. The different indicators present different responses to flooding stress in the study, which may arise from different sensibility of cultivars with different genotypes to flooding stress according to their various physiological parameters and antioxidant system (Do et al., 2018). Moreover, drought stress induces the higher activities of antioxidative enzymes (except APX) and contents of osmotic adjustment substances which promote the growth of maize with a greater LA and plant height rather than flooding stress. Interestingly, the stem diameter in most varieties can become thicker to alleviate the adverse effects of flooding stress relative to drought stress as it is one of the important elements which determines the lodging resistance of maize (Esechie, 1985). Tian et al. (2019) also revealed that the thicker the stem is, the more resistant maize is to flooding stress. Besides, the increment of APX in flooding stress similar to drought stress which is a typical adaptive stress response may play a dominant role in antioxidant indicators. The F_4_ identified through PCA also verified the conclusion. APX activity that controls the H_2_O_2_ scavenging system increased to provide plants with increased flooding stress tolerance which can be used as a standard to evaluate the flooding resistance of plants as well as drought resistance (Lin et al., 2004; Do et al., 2018).

The ROS that are toxic byproducts of oxidative metabolisms are induced the significant increment by drought or flooding stress which are in line with previous studies (Ye et al., 2015; Song et al., 2019), but flooding stress contributes more toward ROS especially O_2_^−^ than drought stress. It can be scavenged through the activated internal protection mechanisms such as the activation of antioxidant enzymes and the production of osmotic adjustment substances (Wang et al., 2019; Jia et al., 2021). Under drought stress, higher activities of antioxidant enzymes and bulk accumulation of osmotic adjustment substances together promote ROS to be scavenged to protect the plants against oxidative damage and maintain ROS to be a lower level. Ye et al. (2015) revealed that oxidative stress occurred in drought but not in submergence stress. However, in the study, maize under flooding suffered from slightly different oxidative stress relative to drought stress. The enzyme activities of SOD and CAT, production of soluble protein, and proline decreased after flooding treatment, which was all identified to be significantly and negatively correlated with O_2_^−^. A previous study found that the lower enzyme activity of SOD was, the less O_2_^−^ converted to H_2_O_2_ was (Keune et al., 2013). Meanwhile, soluble protein and proline also played important roles in the scavenging of ROS. All the above are the reason why more O_2_^−^ is accumulated under flooding stress compared with drought stress. On the other hand, APX and POD catalyze the conversion of H_2_O_2_ to O_2_^−^ (Lin et al., 2004; Salah et al., 2019), leading to the low content of H_2_O_2_ accumulated in both drought and flooding stresses. Moreover, the more accumulation of ROS in flooding stress or drought stress is, the more membrane lipid peroxidation and damage membrane homeostasis are, ultimately accelerating leaf senescence and inhibiting the growth of plants (LIU et al., 2010; Wang et al., 2021). All the above give us new insights into understanding the different scavenging mechanisms of ROS and the adaptive growth mechanism of maize in response to drought and flooding stresses.

The drought and flooding resistances of crops are complex quantitative characteristics which influenced by the combination of genotypes and environment and whose mechanisms are much more complicated. It is a great limit and deficiency to evaluate drought or flooding tolerance only by a single indicator. Meanwhile, overlapping information exists among many evaluation indicators e.g. multiple linear correlations not only reveal the relationships between different indicators but also facilitate the analysis of the drought- or flooding-tolerance mechanism through the internal relationship of indicators. Thus, many scholars have developed different multivariate analysis methods to evaluate drought or flooding tolerance of crops’ comprehensive character index (LIU et al., 2015; Füzy et al., 2019; Shahrokhi et al., 2020). The resistance coefficient whether in drought, flooding, or salt stress has been extensively and generally used in most studies (LIU et al., 2015; Badr et al., 2020; Yu et al., 2021). Although the resistance coefficient well reflects the sensitivity of plants to abiotic stress, that does not reflect the original characteristics of different indexes themselves identified in abiotic stress due to the resemblance of resistance coefficient in most varieties. For example, Guidan 162 identified in the study had the highest drought and flooding resistance coefficients, nevertheless, the contents of most indicators, such as proline, POD, CAT, and LA were lower than those of other varieties when it suffered from drought or flooding stress. Hence, membership function value and PCA are adopted to compensate for the deficiency of resistance coefficient and to evaluate and screen resistant cultivars for abiotic stress (Chen et al. 2012; Bo et al. 2017; Zou et al. 2020). The membership function transforms multiple indexes based on their averages quantitatively which overcomes the one-sidedness of the single indicator, yet it still exists one-sidedness for evaluating the resistance of plants. Instead of the membership function, the D that is calculated by using the membership function and the weight of each index (Yu et al., 2020; Zou et al., 2020) is applied to evaluating the drought and flooding resistances of maize germplasm, which is more comprehensive and accurate than the membership function. It can effectively reflect the comprehensive performance of crops under drought and flooding stresses, and the higher it is, the higher drought- or flooding-resistance capacity of the plant is. Moreover, PCA can simplify the original multiple variables into a few representative indexes namely principal components that are independent to each other (Sun et al., 2021). The important relevant traits for resistance are screened out by PCA and their relationship with each other is revealed by correlation analysis. Furthermore, the key relevant indexes and their specific relationships can be further revealed by stepwise regression analysis. Finally, the maize cultivars are objectively classified through cluster analysis. A multivariate analysis method of PCA, D, correlation analysis, stepwise regression analysis, and cluster analysis was adopted for the study to comprehensively and systematically evaluate and screen the drought and flooding resistances of maize varieties. Four principal components converted from the 10 single indices of maize seedlings measured under drought and flooding treatment by PCA were identified, confirming that antioxidant factor, comprehensive morphological factor, and osmotic adjustment factor all together play a crucial role in improving the tolerance of maize for drought and flooding stresses. The results of correlation analysis also verify the above conclusion as comprehensive principal component (F) is significantly and positively correlated with other indicators. The important and relevant indicators for the comprehensive assessment of drought and flooding stresses of maize varieties are between the D value and the 10 indicators namely soluble protein, sugar, proline, SOD, POD, CAT, APX, LA, plant height, and stem diameter were screened out by stepwise regression analysis those were the same as the results of PCA. According to the optimum regression equations of 12 and 13, all the 10 indicators significantly affect the D value and can determine a relatively high mean forecast accuracy of 99.9% for the D value. Indicators selected in the study also have been more or less screened out in other studies to assess the resistance of plants under various stresses (Bo et at., 2017; Zou et al., 2020; Sun et al., 2021). Consequently, all of the indicators may be much more representative and accurate to be used for the evaluation of drought and flooding resistance of maize varieties at the third-leaves stage.

Due to the importance of D and F values in evaluating the resistance of plants to abiotic stress, and all of them positively and significantly correlated with resistance coefficient, 40 varieties in the present study were successfully clustered into five groups based on the D and F values. A maize variety under different water stress was clustered into different groups due to the different responses to water stress as described above. According to Figure 5, under drought stress, four varieties comprised Guidan 902, Guidan 0810, Guidan 673, and Guidan 670 in group V presented the top four D values and top nine F values, suggesting that these varieties have high drought resistance. Moreover, another six varieties in the group IV with top 10 D values and top 11 F values still exist resistant to drought. Although all the above varieties have top ranks of D and F values, taking into account of drought-resistance coefficient, Guidan 0810 had the highest drought-resistance coefficient, followed by Guidan 673, Guidan 902, and Guidan 670 during 10 varieties. Under flooding stress, varieties of D and F values all ranked top four were clustered into group II involving varieties of Guidan 668, Taipingyang 891, Wanchuan 973, and Guidan 551, in which Guidan 668 had the highest flooding-resistance coefficient of 1.51. Variety of Guidan 668 may be screened as the tolerance to both drought and flooding stresses. Interestingly, Zhaofeng 505 is recognized as the most sensitive cultivar to drought and flooding stresses with the smallest D and F values as well as smaller resistance coefficient to water stress. Overall, a comprehensive analysis method of the D and F values combined with resistance coefficient is suggested to be a reliable and accurate method to evaluate and screen varieties under similar water stress. The 10 indicators selected by PCA and stepwise regression analysis can be used for the identification of maize drought and flooding tolerance and the screening of drought- or flooding-resistant materials.

## Conclusion

In the study, the drought and flooding resistances of 40 maize varieties at the third-leaf stage were compared and determined based on a multivariate analysis method of principal component analysis, comprehensive D value, correlation analysis, stepwise regression analysis, and cluster analysis. Most varieties had stronger drought resistance due to higher antioxidant enzyme activities and osmotic adjustment substances and less ROS compared with flooding resistance leading to the drought-resistance coefficient greater than 1.0. The strikingly different mechanisms from the drought of ROS scavenging and adaptive growth of maize in response to flooding stress were identified. Flooding stress results in the increment of ROS (especially O_2_^−^), APX, POD, and soluble sugar, the decrement of SOD, CAT, soluble protein, and the lower than 1.0 flooding-resistance coefficient in most maize varieties compared with well water. The 10 indices comprised SOD, POD, CAT, APX, proline, soluble sugar, and protein were identified as the much accurate and representative indicators to evaluate the drought and flooding resistance of maize at the seedling stage. Eventually, the drought resistance variety of Guidan 0810, flooding resistance variety of Guidan 668, and water-sensitive variety of Zhaofeng 505 were screened out, and they can be used as ideal experimental materials to further study the mechanism of maize response to water stress.

## Supplementary Data

**The Supplementary Figure S1**. The effects of water stress on soluble protein in leaves of different maize varieties. The bars are standard error.

**The Supplementary Figure S2**. The content of soluble sugar in maize varieties changed with water stress. The bars are standard error.

**The Supplementary Figure S3**. The response of proline in maize varieties to water stress. The bars are standard error.

**The Supplementary Figure S4**. The response of superoxide dismutase (SOD) in maize varieties to water stress. The bars are standard error and the numbers represent the average.

**The Supplementary Figure S5**. The effects of water stress on peroxidase (POD) in maize varieties to water stress. The bars are standard error and the numbers represent the average.

**The Supplementary Figure S6**. The response of catalase (CAT) in maize varieties to water stress. The bars are standard error and the numbers represent the average.

**The Supplementary Figure S7**. The response of ascorbate peroxidase (APX) in maize varieties to water stress. The bars are standard error and the numbers represent the average.

**The SupplementaryFigure S8**. The accumulation of hydrogen peroxide (H_2_O_2_) in leaf of maize in relative to water stress. The numbers represent the average.

**The Supplementary Figure S9**. The accumulation of superoxide anion (O2 ^−^) in leaf of maize in relative to water stress. The numbers represent the average.

**The Supplementary Figure S10**. Loading matrix and variance contribution rate of principal component under water stress. F_1_, F_2_, F_3_, three top principal components; SOD, superoxide dismutase; POD, peroxidase; CAT, catalase; APX, anticyclohexate peroxidase

**The Supplementary Table S1**. Morphological indexes in maize under water stress.

### Funding

This work was supported by the Innovation Project of Guangxi Graduate Education (YCBZ2022001) and Science and Technology Development Fund of Guangxi Academy of Agricultural Sciences (GNK2019ZX13).

## Acknowledgments

We thank Dr. Haiyu Zhou for providing some experimental material.

